# Secreted Frizzled-Related Protein 1a regulates hematopoietic development in a dose-dependent manner

**DOI:** 10.1101/2025.01.10.632371

**Authors:** Amber D. Ide, Kelsey A. Carpenter, Mohamed Elaswad, Katherine Opria, Kendersley Marcellin, Carla Gilliland, Stephanie Grainger

## Abstract

Hematopoietic stem and progenitor cells (HSPCs) arise only during embryonic development, and their identity specification, emergence from the floor of the dorsal aorta, and proliferation are all tightly regulated by molecular mechanisms such as signaling cues. Among these, Wnt signaling plays an important role in HSPC specification, differentiation, and self-renewal, requiring precise modulation for proper development and homeostasis. Wnt signaling is initiated when a Wnt ligand binds to cell surface receptors such as those encoded by the *frizzled* gene family, activating intracellular signaling pathways that regulate gene expression. Secreted frizzled-related proteins (Sfrps) are known modulators of Wnt signaling, acting as both agonists and antagonists of this pathway. Yet, *in vivo* functions of Sfrps in HSPC development remain incompletely understood. Here, we demonstrate that Sfrp1a regulates zebrafish HSPC development and differentiation in a dose-dependent manner. In Sfrp1a loss of function animals, we observe an increase in HSPCs, an upregulation of canonical Wnt signaling, and a decrease in differentiation into both lymphoid and myeloid lineages. Conversely, at low-dose *sfrp1a* overexpression, there is a decrease in HSPCs and an increase in lymphoid differentiation. High-dose *sfrp1a* overexpression phenocopies the loss of function animals, with an increase in HSPCs, increased canonical Wnt signaling, and decreased lymphoid and myeloid differentiation. These findings highlight the importance of dose-dependent modulation of Sfrps, paralleling what is observed in hematopoietic cancers where SFRP1 loss-of-function and gain-of-function variants can drive tumorigenesis.

**One sentence summary:** Sfrp1a is required for hematopoietic stem cell development.

## INTRODUCTION

Hematopoietic stem cells (HSCs) are multipotent stem cells essential for generating all blood cell lineages throughout an organism’s life^1,2^. During embryonic development, HSCs originate from hemogenic endothelium, a specialized group of endothelial cells derived from endothelial progenitors^3,4,5^. Nearby tissues, such as the somite and neural crest cells, provide signaling cues that dictate identity specification, emergence and function of hematopoietic stem and progenitor cells (HSPCs)^3,6,7,8,9^. After emergence, HSPCs enter circulation, migrating to intermediate hematopoietic sites like the fetal liver in mammals or the caudal hematopoietic tissue in teleost, before establishing their long-term residence in the bone marrow or the kidney marrow of teleosts^10,11^.

The Wnt signaling pathway is a critical regulator of HSPC development, maintenance, and homeostasis^12,13,14,15,16,17,18,19,20,21,22,23,24,25^. The canonical, or β-catenin-dependent, Wnt signaling pathway is initiated when a Wnt ligand binds to a Frizzled (Fzd) receptor and co-receptors such as the low-density lipoprotein receptor-related protein 5/6 on the cell surface^26,27,28,29,30,31,32^. These interactions activate intracellular signaling cascades that stabilize β-catenin, allowing it to accumulate in the cytoplasm and translocate into the nucleus^29,31,33,34,35^. In the nucleus, β-catenin interacts with transcriptional regulators, including members of the T-cell factor/lymphoid enhancer factor family, to modulate gene expression programs critical for HSPC self-renewal, proliferation, and differentiation^16,36,37,38,39^. Due to its importance in these processes, the level of Wnt signaling output must also be tightly regulated.

The concept of a “just-right” level of canonical Wnt signaling has emerged as an essential mechanism for HSPC function and deviations from this can lead to hematological malignancies^15,40^. Although we now understand some of the ligands and receptors required for HSPC development, the modulators that fine tune signaling levels are incompletely understood. Among these modulators are Secreted Frizzled-Related Proteins (SFRPs), a family of extracellular proteins capable of interacting with Wnt ligands and/or Fzd receptors. In humans, five SFRP proteins have been identified, and orthologs of these genes are present in all vertebrate species^41^. Each SFRP protein contains a cysteine-rich domain, similar to the ligand binding domain of Fzd receptors, and a netrin-related domain^42,43,44^. Both loss- and gain-of-function of SFRPs, particularly SFRP1, have been implicated in various cancers, including those of the blood^45,45,46,47,48^. While traditionally considered as Wnt inhibitors, emerging evidence suggests that SFRPs can also function as activators of Wnt signaling^49,50,51,52,53^. These bimodal functions may be related to dose-dependent differences in function^53^, although this is less clear *in vivo*.

Here, we sought to investigate Sfrp regulation of HSPC development *in vivo* using zebrafish as a model system. We have identified Sfrp1a as a critical regulator of HSPC development and blood cell differentiation. Consistent with findings from *in vitro* studies, our findings demonstrate a dose-dependent role of Sfrp1a *in vivo*, emphasizing the importance of understanding how proteins within the Wnt signaling pathway operate across varying concentrations.

## RESULTS

### Sfrp1a is expressed in endothelial cells and influences HSPC development

In zebrafish, there is a critical time window for both canonical and non-canonical Wnt signaling prior to 20 hours post fertilization (hpf)^6,16,17^. The Sfrp(s) modulating Wnt function in HSPC development remain unexplored. We hypothesized that Sfrps, known to modulate Wnt signaling, may be expressed in HSPC-adjacent tissues, such as endothelial cells, during this developmental window. To investigate which zebrafish *sfrp(s)* are expressed during the window of Wnt-driven HSPC development, we performed qPCR on endothelial cells at 16.5 hpf from *fli1a:eGFP* zebrafish (Fig. 1A) and found that *sfrp1a* was the most highly expressed at this stage (Fig. 1B). Whole-mount *in situ* hybridization (WISH) confirmed *sfrp1a* expression in endothelial cells at 13, 16.5 and 19 hpf (Fig. 1C), during the Wnt-responsive HSPC developmental window.

**Figure 1:**
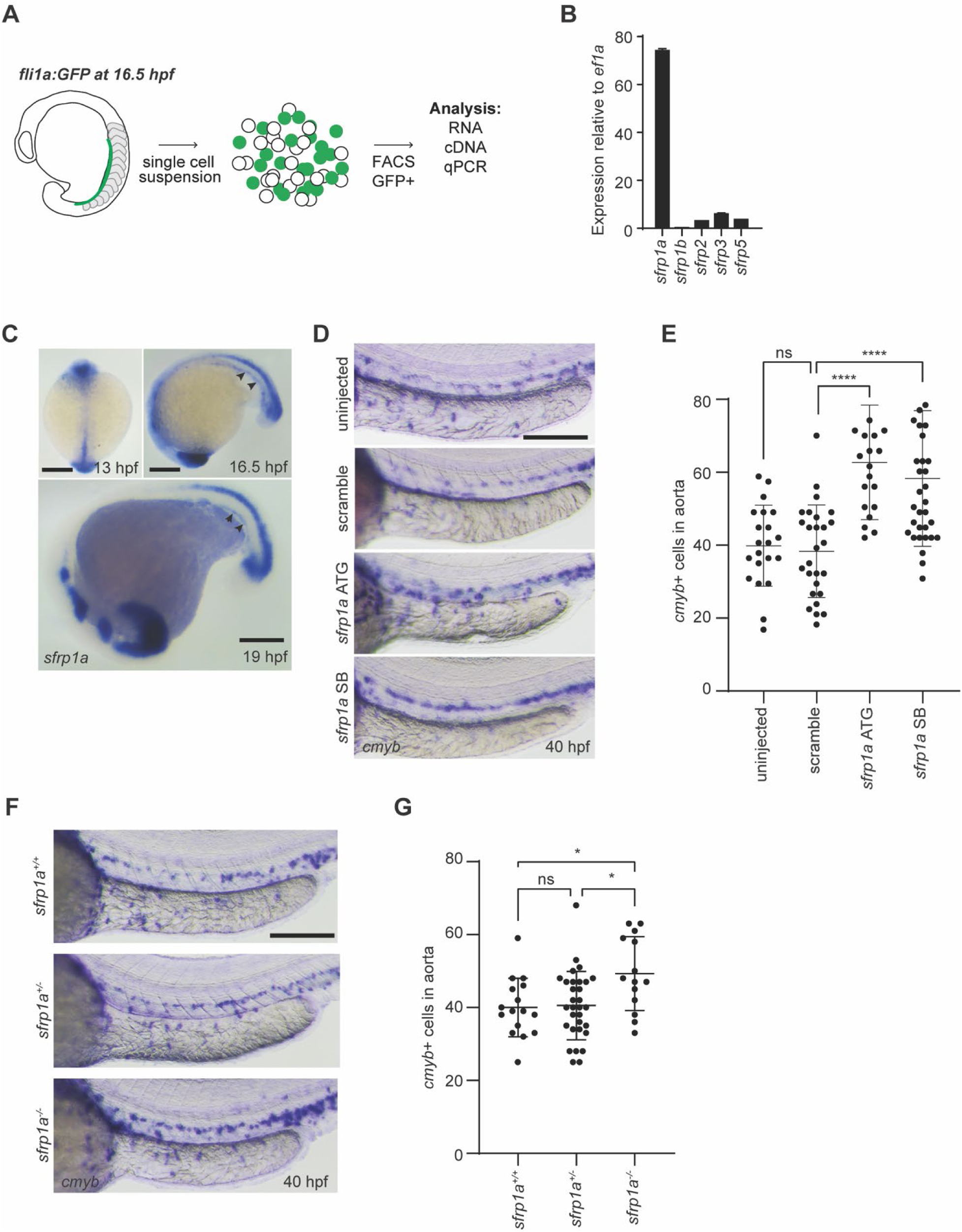
Sfrp1a is expressed in endothelial cells and affects HSPC development. **A**. *fli1a:GFP* fish were dissociated at 16.5 hpf and endothelial cells (GFP+) were collected by FACS for qPCR. **B.** qPCR for each zebrafish *sfrp* (*n*= 3 biological replicates). **C.** Representative WISH images for *sfrp1a* at 13 hpf (scale bar = 30 µm), 16.5 hpf (scale bar = 30 µm), and 19 hpf (scale bar = 50 µm). Arrow heads indicating *sfrp1a* expression in endothelial cells. **D.** AB* embryos were injected with scramble morpholino and *sfrp1a* morpholinos: ATG and splice blocking (SB), fixed at 40 hpf, and analyzed by WISH for *cmyb*. Arrow heads in representative images point at HSPCs in the floor of the dorsal aorta. Scale bar = 100 µm. **E.** Quantification of **D** (n=21, 27, 20, and 34 biological replicates from left to right). Each dot represents a biological replicate, the bars represent the mean and the error bars represent the standard deviation. One-Way ANOVA with post hoc Tukey comparisons, **** p<0.0001. **F.** *sfrp1a* mutants were generated by CRISPR/Cas9 and representative WISH images for *cmyb* expression at 40 hpf. Scale bar = 100 µm. **G.** Quantification of **F** (*n*= 16, 31, and 14 biological replicates from left to right). Each dot represents a biological replicate, the bars represent the mean and the error bars represent the standard deviation. One-Way ANOVA with post hoc Tukey comparisons, * p<0.05.

HSPC development can be assessed by examining *cmyb+* cells in the dorsal aorta at 40 hpf^16^. To determine if Sfrp1a is required for HSPC development, we used an ATG and splice-blocking (SB) morpholinos (MO) to knock down *sfrp1a* expression during development. Knockdown of *sfrp1a* resulted in a significant increase in the number of *cmyb*+ HSPCs in the aorta (Fig. 1D,E). We also noted a 3-fold increase in *cmyb* expression at 40 hpf by qPCR (Fig. S1A), suggesting that Sfrp1a influences HSPC development. To confirm specificity of the Sfrp1a MOs, we used CRISPR/Cas9-mediated mutagenesis to target the proximal promoter and translational start site of the *sfrp1a* coding region, which resulted in a 515 bp deletion (Fig. S1B) predicted to result in a null allele (hereafter referred to as *sfrp1a^-/-^*). These *sfrp1a^-/-^* animals also displayed a significant increase in *cmyb+* HSPCs in the aorta (Fig. 1F, G), supporting the specificity of the MO. In addition, *sfrp1a* MO injection did not impact the phenotype of *sfrp1a^-/-^* animals (Fig. S1C), further confirming that our MO and mutant animals exhibit HSPC phenotypes due to the loss of HSPCs since it did not disrupt the development of primitive blood (*gata1a*), somites (*myod*), or the vasculature (*kdrl*) (Fig. S1D). HSPC emergence begins around 26 hpf and is marked by the appearance of *runx1+* HSPCs in the dorsal aorta, a process that is dependent upon non-canonical Wnt signaling^54^. In *sfrp1a* MO-injected embryos, no defects were observed in the number of *runx1+* cells at 26 hpf (Fig. S1E), indicating that Sfrp1a does not impact HSPC emergence.

### *sfrp1a* expression prior to HSPC emergence influences HSPC development

Since *sfrp1a* knock down increased HSPCs at 40 hpf, we hypothesized that overexpressing *sfrp1a* would decrease HSPCs at 40 hpf. We injected *sfrp1a* mRNA into zebrafish embryos and analyzed *cmyb* expression at 40 hpf using WISH and qPCR. As expected, with a 50 pg dose of *sfrp1a* mRNA, we observed a significant decrease in *cmyb+* HSPCs in the aorta at 40 hpf (Fig. 2A), which we confirmed by qPCR (Fig. 2B).

**Figure 2:**
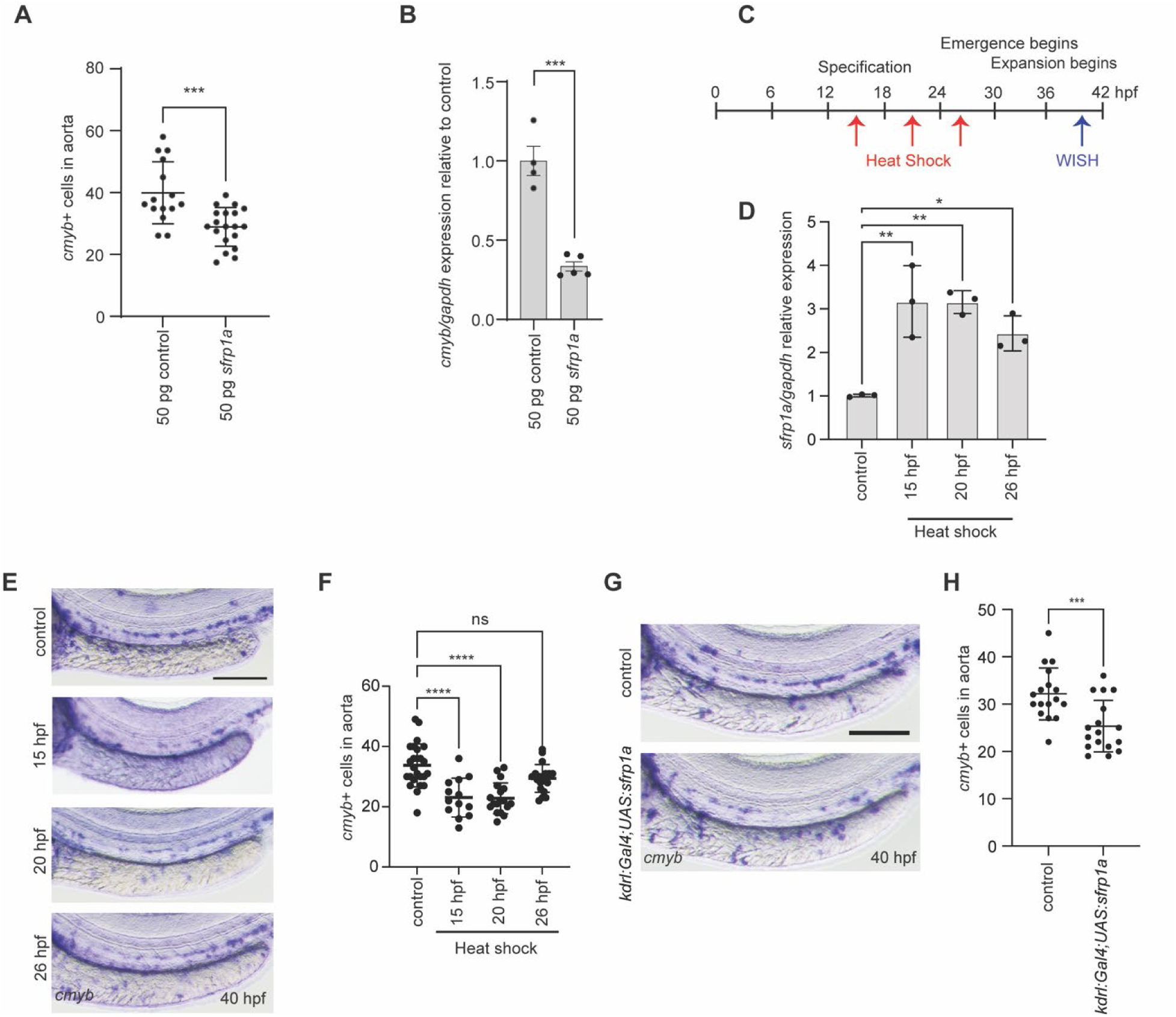
*sfrp1a* expression prior to HSPC emergence influences HSPC development. **A.** AB* embryos were injected with 50 pg control mRNA or 50 pg *sfrp1a* mRNA, fixed at 40 hpf, and analyzed by WISH for *cmyb* (*n*= 15 and 19 biological replicates from left to right). Each dot represents a biological replicate, the bars represent the mean and the error bars represent the standard deviation. One-Way ANOVA with post hoc Tukey comparisons, ***p<0.001. **B.** AB* embryos were injected with 50 pg control mRNA or 50 pg *sfrp1a* mRNA and expression of *cmyb* was analyzed using qPCR (*n=* 5 fish for each biological replicate, 3 biological replicates). Two-tailed Student’s t test, ***p<0.001. **C.** Schematic of heat shock regimen during important HSPC developmental time windows. **D.** *hsp:sfrp1a* fish were heat shocked for 30 minutes at 35°C, allowed to recover to 28.5°C for an hour and then fish were collected for *sfrp1a* qPCR (*n=* 5 fish for each biological replicate, 3 biological replicates). One-Way ANOVA with post hoc Tukey comparisons, *p<0.05, **p<0.01. **E.** Representative images of *hsp:sfrp1a* fish that were heat shocked at 15 hpf, 20 hpf, or 26 hpf for 30 minutes at 35°C, fixed at 40 hpf, and analyzed by WISH for *cmyb*. **F.** Quantification of **E** (n= 25, 13, 18, 19 biological replicates from left to right). Each dot represents a biological replicate, the bars represent the mean and the error bars represent the standard deviation. One-Way ANOVA with post hoc Tukey comparisons, ****p<0.0001.**G.** Representative images of *kdrl:gal4* fish that were crossed with *UAS:sfrp1a* fish, fixed at 40 hpf, and analyzed by WISH for *cmyb*. Control fish are non-transgenic siblings with identical heat shock conditions. Scale bar = 100 µm. **H.** Quantification of **G** (*n*= 17 biological replicates). Each dot represents a biological replicate, the bars represent the mean and the error bars represent the standard deviation. Two-tailed Student’s t test, ***p<0.001.

To explore the spatial and temporal requirements for Sfrp1a during HSPC development, we generated *UAS:sfrp1a* transgenic animals, which can be induced in specific tissues by *Gal4* drivers. To test that Gal4 induced expression of the *sfrp1a* transgene, we generated *hsp70l:Gal4;UAS:sfrp1a* animals and performed heat shocks during the Wnt-responsive HSPC development window (15 hpf), just after the requirement for canonical Wnt signaling (20 hpf), and during early HSPC emergence (26 hpf) (Fig. 2C). Heat shocking at all timepoints led to an increase in *sfrp1a* expression (Fig. 2D), indicating that our transgene was functional. Heat shock at the 20 hpf timepoint resulted in a significant decrease in *cmyb+* HSPCs in the aorta at 40 hpf, while heat shocks at 15 hpf and 26 hpf had no effect (Fig. 2E, F), suggesting that Sfrp1a is required prior to HSPC emergence to influence HSPC development.

Sfrp1a is expressed in endothelial cells during the Wnt-responsive HSPC development window prior to 20 hpf; however, neighboring tissues such as the somites also influence HSPC window prior to 20 hpf; however, neighboring tissues such as the somites also influence HSPC development at this stage. To determine if Sfrp1a function in HSPC ontogeny requires tissue-specific expression, we combined *UAS:sfrp1a* animals with several tissue-specific *Gal4* drivers. Like global mRNA overexpression (Fig. 2A, B), exogenous expression of Sfrp1a in all endothelial cells (using *kdrl:Gal4*) led to a significant decrease in *cmyb+* HSPCs at 40 hpf (Fig. 2G, H), indicating that *sfrp1a* overexpression in endothelial cells alone can modulate HSPC development. The canonical Wnt signal is received by cells of the hemogenic endothelium^16^. To assess if overexpression of *sfrp1a* from hemogenic endothelium (driven by *gata2b* promoter regions) is sufficient to alter HSPC development, we examined *cmyb+* cells at 40 hpf in *gata2b:KalTA4; UAS:sfrp1a* animals. We did not observe a difference in *cmyb+* HSPCs at 40 hpf (Fig. S2A), suggesting that *sfrp1a* expression in the hemogenic endothelium does not affect HSPC development. Other neighboring tissues, such as the somites, are known to influence HSPC development^6,55^. To assess if exogenous expression of *sfrp1a* from the somites can alter HSPC development, we examined *phldb1:Gal4;UAS:sfrp1a* animals and did not observed a difference in *cmyb+* HSPCs at 40 hpf (Fig. S2B), indicating that *sfrp1a* expression in somites is not able to alter HSPC development. Taken together, these data suggest a proximity requirement for Sfrp1a to function in HSPC development.

### Sfrp1a loss of function leads to increased Wnt signaling in the developing endothelium

Since loss of Wnt signaling during development leads to a loss of HSPCs at 40 hpf^16^ and Sfrps are proposed to function as antagonists in Wnt signaling, we hypothesized that *sfrp1a* loss of function would lead to an increase in Wnt signaling. To assess how the loss of *sfrp1a* affects canonical Wnt signaling in the developing endothelium, we injected *sfrp1a* ATG MO into *7XTCF:eGFP;kdrl:mCherryNLS* animals. In these animals, GFP is expressed under the control of a Wnt responsive promoter (*7XTCF*)^56^ and nuclear-localized mCherry marks endothelial cells^4^. Consistent with antagonist function, we observed a significant increase in GFP fluorescence in endothelial cells in *sfrp1a* morphant animals during the Wnt-responsive window at 16.5 hpf (Fig. 3A, B). To evaluate the impact of *sfrp1a* overexpression on Wnt signaling at this timepoint, we injected 50 pg of *sfrp1a* mRNA into *7XTCF:eGFP;kdrl:mCherryNLS* embryos and assessed GFP fluorescence in endothelial cells. We did not observe a noticeable difference in GFP fluorescence compared to the control mRNA (Fig. S3A, B). However, the *sfrp1a* loss of function phenotype may suggest that Sfrp1a may antagonize Wnt signaling during HSPC ontogeny.

**Figure 3:**
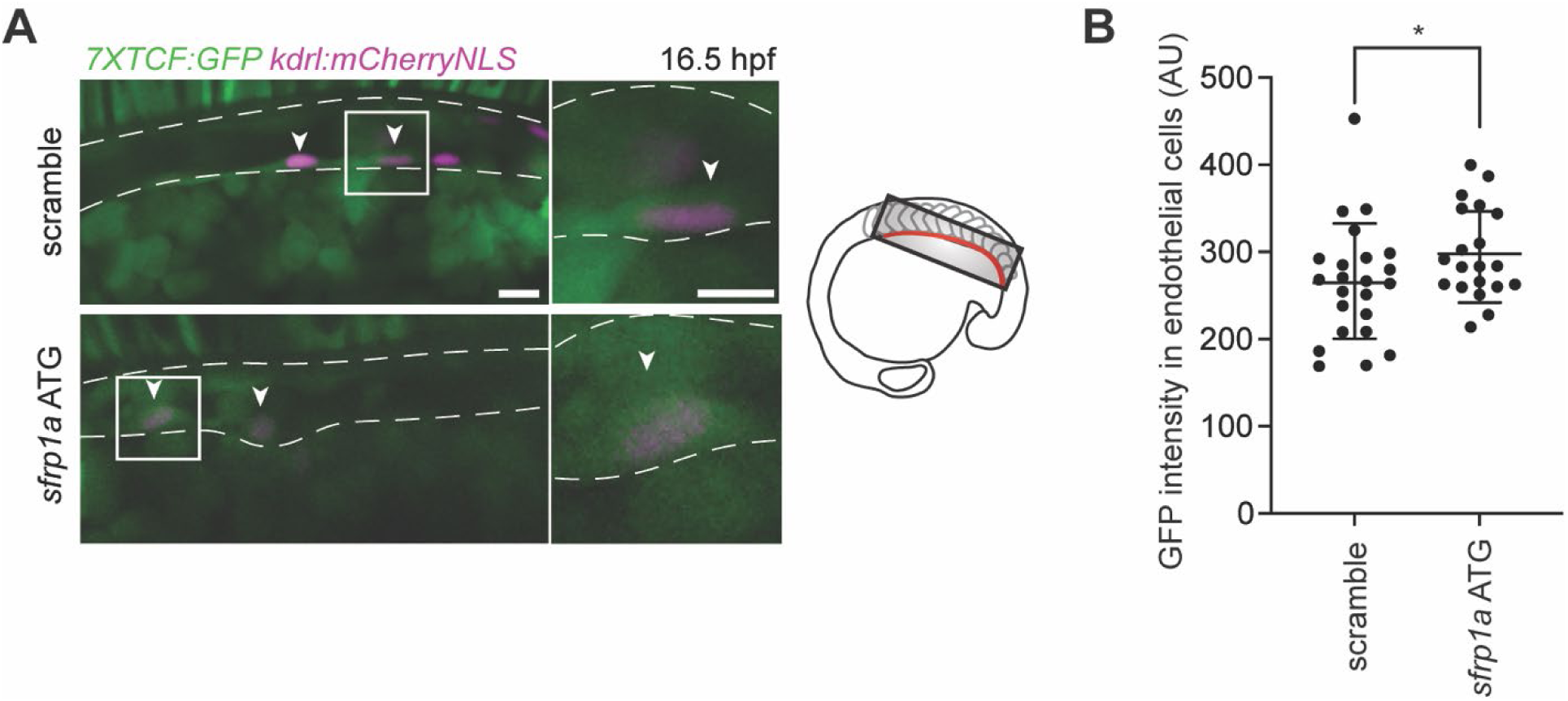
Sfrp1a loss of function leads to increased Wnt signaling in the developing endothelium. **A.** Representative images of *7XTCF:GFP; kdrl:mCherryNLS* fish injected with scramble or *sfrp1a* ATG morpholino and the double positive cells were analyzed for GFP fluorescence at 16.5 hpf. Arrow heads point to double positive endothelial cells. Scale bars= 40 µm and inset is 10 µm. **B.** Quantification of GFP intensity in endothelial cells (*n*= 23 and 20 cells in 9 fish). Two-Way ANOVA, *p<0.05.

### Sfrp1a regulates HSPC development and differentiation in a dose-dependent manner

*In vitro* studies have revealed that SFRPs regulate Wnt ligand delivery to cellular targets in a dose-dependent manner^53^. To test the impact of high levels of Sfrp1a, we injected 500 pg of *sfrp1a* mRNA and observed the opposite effect compared to the 50 pg *sfrp1a* overexpression: a significant increase in *cmyb+* HSPCs in the aorta (Fig. 4A) and increased *cmyb* expression (Fig. 4B) at 40 hpf. We injected 500 pg *sfrp1a* mRNA into *7XTCF:eGFP;kdrl:mCherryNLS* embryos and observed a significant increase in GFP fluorescence in endothelial cells at 16.5 hpf (Fig. S3C, D), similar to *sfrp1a* loss of function. These data confirmed that Sfrp1a function in a dose-dependent manner *in vivo*, similar to *in vitro* findings.

**Figure 4:**
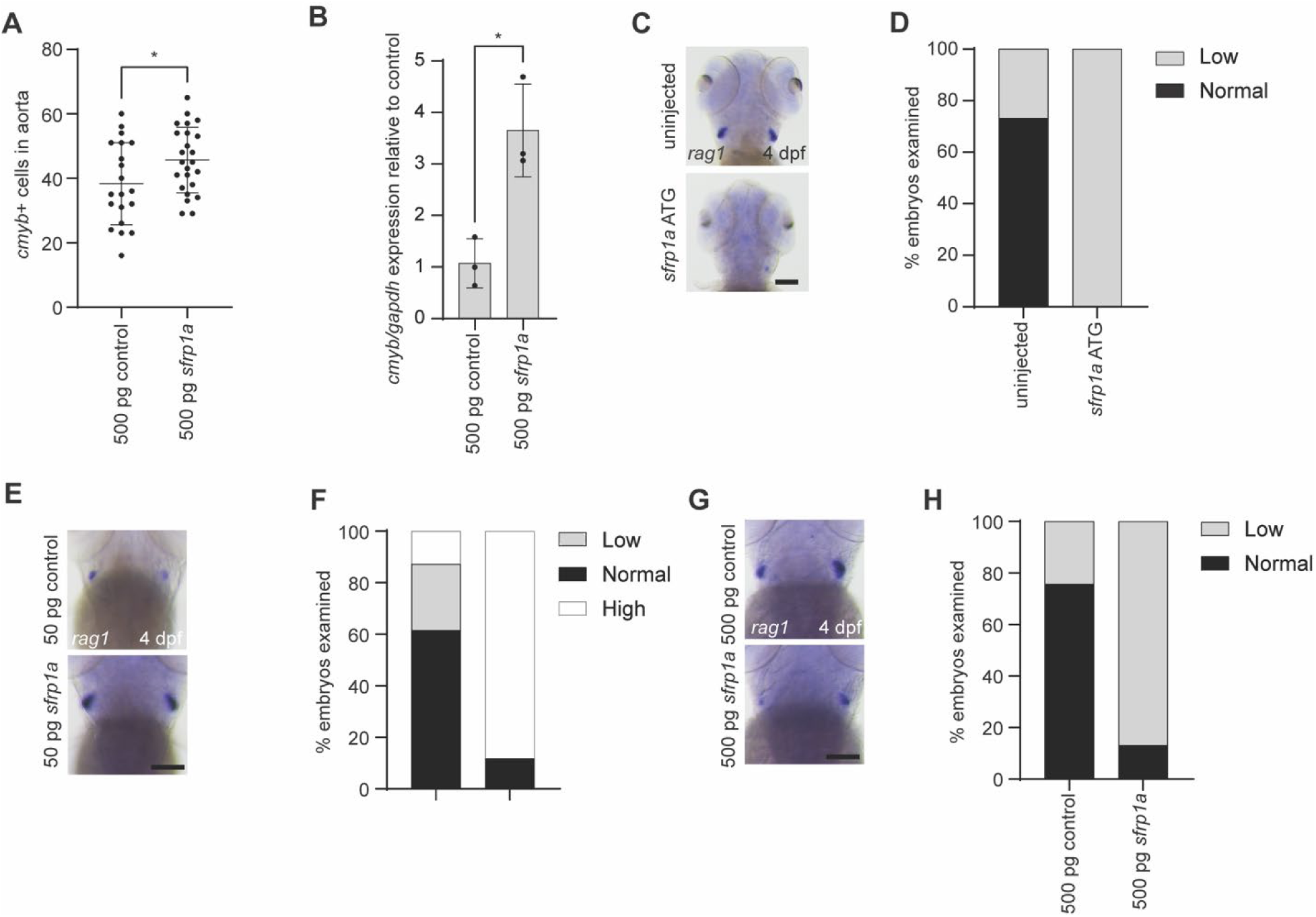
Sfrp1a impacts lymphoid differentiation in a dose-dependent manner. **A.** AB* embryos were injected with 500 pg control mRNA or 500 pg *sfrp1a* mRNA, fixed at 40 hpf, and analyzed by WISH for *cmyb* (*n*= 20 and 24 biological replicates from left to right). Each dot represents a biological replicate, the bars represent the mean and the error bars represent the standard deviation. One-Way ANOVA with post hoc Tukey comparisons, *p<0.05. **B.** AB* embryos were injected with 500 pg control mRNA or 500 pg *sfrp1a* mRNA and expression of *cmyb* was analyzed using qPCR (*n=* 5 fish for each biological replicate, 3 biological replicates). Two-tailed Student’s t test, *p<0.05. **C.** Representative images of uninjected AB* embryos or injected *sfrp1a* ATG morpholino embryos, fixed at 4 dpf, and analyzed by WISH for *rag1.* Scale bar = 50 µm. **D.** Quantification of **C** (*n*= 22 and 26 biological replicates from left to right). **E.** Representative images of AB* embryos injected with 50 pg control mRNA or 50 pg *sfrp1a* mRNA, fixed at 4 dpf, and analyzed by WISH for *rag1.* Scale bar = 50 µm. **F.** Quantification of **E** (*n*= 39 and 34 biological replicates from left to right). **G.** Representative images of AB* embryos injected with 500 pg control mRNA or 50-pg *sfrp1a* mRNA, fixed at 4 dpf, and analyzed by WISH for *rag1.* Scale bar = 50 µm. **H.** Quantification of **G** (*n*= 37 and 38 biological replicates from left to right).

In addition to its roles in HSPC specification and self-renewal, Wnt signaling plays an important role in blood cell differentiation by maintaining the balance between HSPCs and their differentiation into erythro-myeloid progenitors and cells (such as macrophages, neutrophils, monocytes, and erythrocytes) or lymphoid progenitors and cells (such as T cells, B cells, and natural killer cells)^15,57^. To investigate whether Sfrp1a dosage influences the differentiation of HSPCs into lymphoid cells, we injected embryos with *sfrp1a* MO, *sfrp1a* low (50 pg) and high (500 pg) overexpression doses and assessed the formation of thymocytes, the precursors to T cells. We found that *sfrp1a* MO animals had fewer *rag1+* thymocytes (Fig. 4C, D) and expression (Fig. S4A) at 4dpf. Interestingly, this effect was dose-dependent: low doses of *sfrp1a* mRNA increased *rag1+* thymocytes (Fig. 4E, F) and expression (Fig. S4B), whereas high doses caused a decrease in *rag1+* thymocytes (Fig. 4G, H) and expression (Fig. S4C) and 4 dpf. These results demonstrate that Sfrp1a regulates the differentiation into lymphoid lineages in a dose-dependent manner.

To investigate the impact of Sfrp1a loss- and gain-of-function on erythro-myeloid lineages, we conducted similar experiments using *mpeg1:eGFP* transgenic animals, which express GFP in macrophages. Similar to our findings in thymocytes, *sfrp1a* MO-injected animals had fewer *mpeg:eGFP+* cells (Fig. 5A, B); however, we did not observe any differences in *mpeg+* cells at either low dose (Fig. 5C, D) or high dose *sfrp1a* overexpression (Fig. 5E, F). We also observed a decrease in neutrophils, another cell downstream of myeloid progenitors, marked by *mpx* expression, in *sfrp1a* morphant animals (Fig. 5G, H). Interestingly, we did not observe any difference in neutrophils in response to the 50 pg *sfrp1a* overexpression (Fig. 5I, J), but there was a decrease in neutrophils in the 500 pg *sfrp1a* overexpression (Fig. 5K, L). We conducted similar analyses in *gata1:dsRed* animals, which labels erythrocytes, and did not note any changes in response to *sfrp1a* MO or 50 pg *sfrp1a* overexpression (Fig. S5A-D). However, we did see a significant decrease in *dsRed+* cells at the 500 pg overexpression (Fig. S5E, F). These data suggests that Sfrp1a may not impact erythro-myeloid lineage differentiation in a dose-dependent manner similar to lymphoid lineages. However, these results indicate that Sfrp1a plays a role in both lymphoid and myeloid cell lineage differentiation.

**Figure 5.**
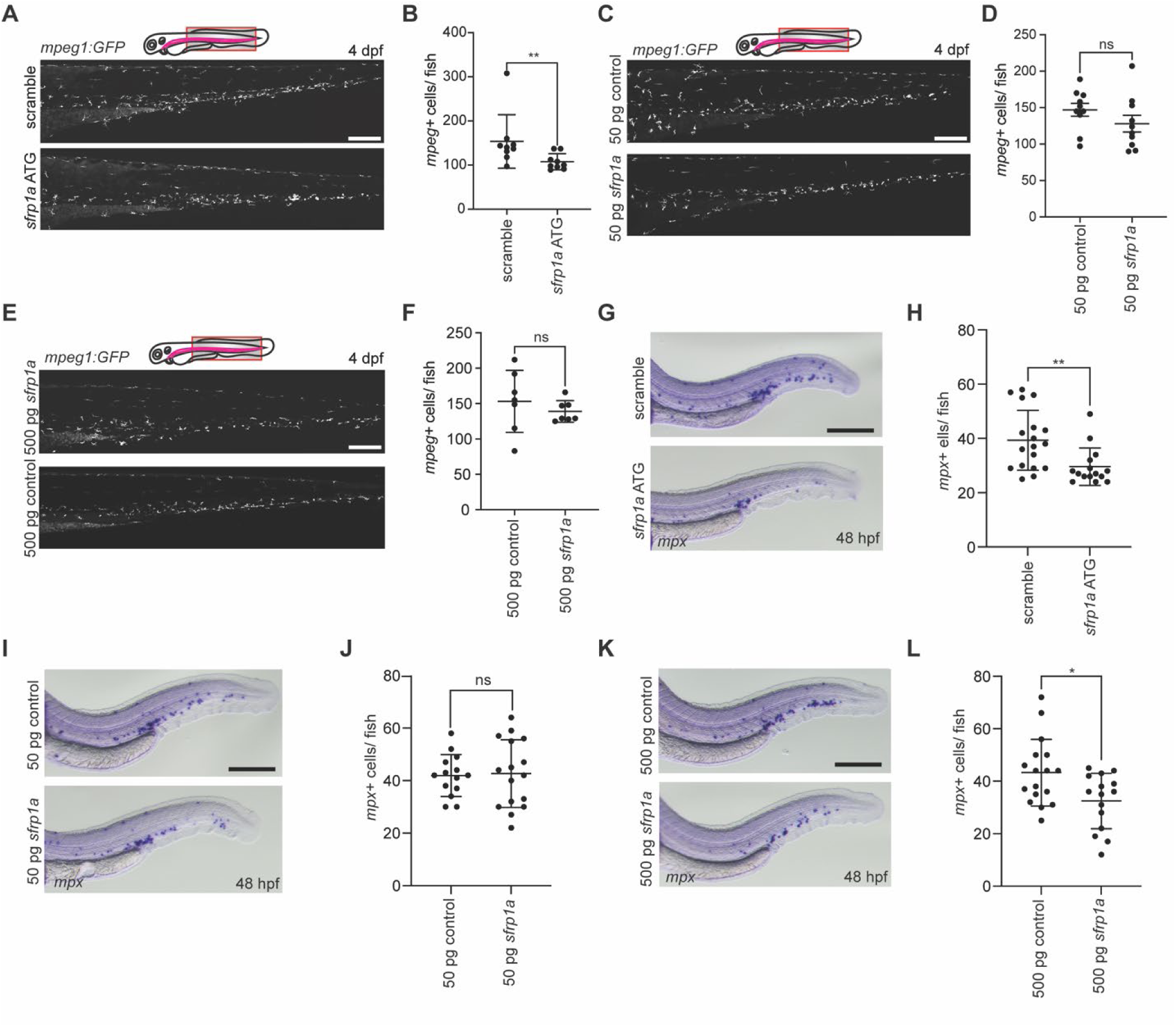
Sfrp1a impacts myeloid differentiation. **A.** Representative confocal Z-stacks of *mpeg1:GFP* fish injected with scramble or *sfrp1a* ATG morpholino, embedded in agar at 4 dpf and imaged at 25X. Scale bar = 200 µm. **B**. Quantification of **A** marked by GFP+ cells (*n*= 9 biological replicates). Each dot represents the mean number of cells in a biological replicate, the bars represent the mean and the error bars represent the standard deviation. Mann-Whitney test, **p<0.01. **C**. Representative confocal Z-stacks of *mpeg1:GFP* fish injected with 50 pg control mRNA or 50 pg *sfrp1a* mRNA, embedded in agar at 4 dpf and imaged at 25X. Scale bar = 200 µm. **D**. Quantification of **C** marked by GFP+ cells (*n*= 10 biological replicates). Each dot represents the mean number of cells in a biological replicate, the bars represent the mean and the error bars represent the standard deviation. Mann-Whitney test. **E**. Representative confocal Z-stacks of *mpeg1:GFP* fish injected with 500 pg control mRNA or 500 pg *sfrp1a* mRNA, embedded in agar at 4 dpf and imaged at 25X. Scale bar = 200 µm. **F**. Quantification of **E** marked by GFP+ cells (*n*= 7 biological replicates). Each dot represents the mean number of cells in a biological replicate, the bars represent the mean and the error bars represent the standard deviation. Mann-Whitney test. **G.** Representative images of embryos injected with scramble or *sfrp1a* ATG morpholino, fixed at 48 hpf, and analyzed by WISH for *mpx.* Scale bar = 125 µm. **H.** Quantification of **G** (*n*= 18 and 15 biological replicates from left to right). Each dot represents the mean number of cells in a biological replicate, the bars represent the mean and the error bars represent the standard deviation. Two-tailed student’s t-test, **p<0.01. **I.** Representative images of embryos injected with 50 pg control mRNA or 50 pg *sfrp1a* mRNA, fixed at 48 hpf, and analyzed by WISH for *mpx.* Scale bar = 125 µm. **J.** Quantification of **I** (*n*= 14 and 16 biological replicates from left to right). Each dot represents the mean number of cells in a biological replicate, the bars represent the mean and the error bars represent the standard deviation. Two-tailed student’s t-test. **K.** Representative images of embryos injected with 500 pg control mRNA or 500 pg *sfrp1a* mRNA, fixed at 48 hpf, and analyzed by WISH for *mpx.* Scale bar = 125 µm. **L.** Quantification of **K** (*n*= 17 and 15 biological replicates from left to right). Each dot represents the mean number of cells in a biological replicate, the bars represent the mean and the error bars represent the standard deviation. Two-tailed student’s t-test, *p<0.05.

## DISCUSSION

Wnt signaling is critical for HSPC development and homeostasis. However, the role of pathway modulators like Sfrps during HSPC development remains incompletely understood. Sfrp proteins can function both as agonists and antagonists in a dose-dependent manner *in vitro*, adding complexity to their study and functional characterization. Here, we investigated the role of Sfrps during HSPC development in zebrafish.

Our findings present the first study to demonstrate that Sfrp1a plays an important role in HSPC development, acting as a key regulator of HSPC differentiation in a dose-dependent manner *in vivo*. Sfrp1a loss of function and overexpression resulted in differences in HSPC numbers due to impaired HSPC differentiation into lymphoid and myeloid lineages (Fig. 6A). These results support previous *in vitro* studies demonstrating dose-dependent effects of Sfrps and highlight the role of Sfrp1a in HSPC differentiation into lymphoid and myeloid lineages (Fig. 6B).

**Figure 6:**
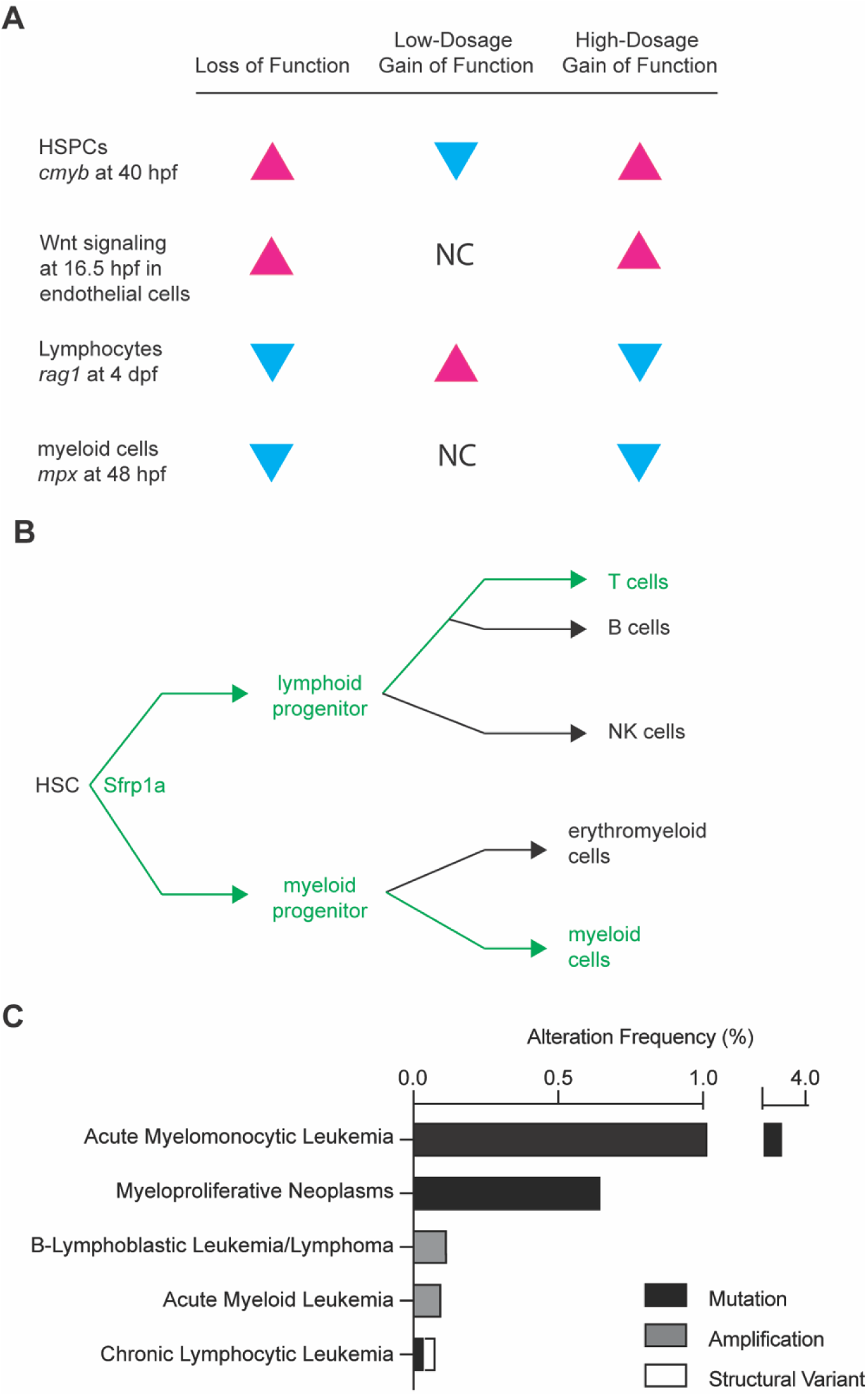
Sfrp1a is required for proper HSPC development and differentiation. **A.** Table summarizing main findings. Magenta arrow heads indicate an increase while the blue arrow heads indicate a decrease. NC indicates no change. **B.** Schematic of where Sfrp1a may regulate HSPC differentiation into lymphoid and myeloid lineages. **C.** Alteration frequency of SFRP1variants in human hematopoietic malignancies^72,73,74^.

Traditionally, Sfrp proteins were though to either deliver Wnt ligands to Fzd receptors or sequester them away to inhibit signaling; however, emerging evidence suggests bimodal functions at different doses^49,50,51,52,53^. SFRPs bind Wnt ligands and facilitate their transport to glypican proteoglycans (GPC) on the cell surface, which act as docking sites for Wnt ligands^53^. At low SFRP concentrations *in vitro*, SFRPs promote the efficient transfer of Wnt ligands from GPCs to Fzd receptors, thereby enhancing Wnt signaling. In contrast, at high concentrations, SFRPs sequester Wnt ligands, reducing their availability to Fzd receptors and attenuating Wnt signaling^53^. While these findings provide significant insights into SFRP-mediated modulation of Wnt signaling, the dose-dependent mechanism has yet to be confirmed *in vivo.* Our investigations of Sfrp1a function are in line with this model. In *sfrp1a* loss of function models, there is an increase in Wnt signaling in the endothelium, leading to increased HSPC development (Fig. 1, 3), and impaired differentiation (Fig. 4, 5), in line with Sfrp1a acting as an antagonist normally, and loss of function leading to relief of repression. When *sfrp1a* is overexpressed at a low dose, we also observed this function as an antagonist, with decreased HSPC development (Fig. 2) and increased lymphoid differentiation (Fig. 4). Interestingly, when we exogenously expressed a high dose of *sfrp1a*, we observed an increase in endothelial Wnt signaling (Fig. S3C, D), concomitant with an increase in HSPC development (Fig. 4) and decreased lymphoid and myeloid differentiation (Fig. 4, 5). This could be due to Sfrp1a directly binding to Fzd receptors and competitively inhibiting Wnt ligand interactions, a phenomenon observed in overexpression studies of SFRPs *in vitro*^53^. Future studies exploring the interaction of Sfrp1a with GPCs and Fzd receptors *in vivo* are needed to validate this model.

We observed an increase in Wnt signaling in the *sfrp1a* loss of function and high dose models; however, we did not observe a decrease in Wnt signaling at low *sfrp1a* overexpression. We did, nevertheless, observe a phenotype consistent with decreased Wnt signaling. This inconsistency may reflect (1) not examining the appropriate cell type or developmental timepoint where Sfrp1a at this dose impacts canonical Wnt signaling, or (2) that there are multiple Wnt pathways at play, and the changes in signaling intensity with Sfrp1a modulation are minor comparatively. This observation is consistent with our observations that Wnt9a/Fzd9b signaling, which is critical to HSPC development, does not induce *in vitro* Wnt reporter to the same degree as Wnt3a^16,17^. Furthermore, we observed a clear dose-dependent differentiation response in lymphoid lineages whereas this pattern was less apparent in the myeloid lineage. This discrepancy could suggest differing dose requirements between these cell types, the possibility of having missed the optimal timepoint for observation, or that Sfrp1a may not regulate myeloid differentiation in a dose-dependent manner. This just further highlights that Sfrp1a plays a more complex role *in vivo* than *in vitro* studies indicate. Others have also described the context-dependent functions for different Wnt pathways and other influences such as cell-type specificity, co-factors, or variations in the local Wnt signaling environment^58,59,60^. The spatiotemporal expression of Wnt ligands, Fzd receptors, and Sfrps further defines Wnt signaling specificity. For example, Wnt9a/Fzd9b interactions are required for HSPC proliferation in zebrafish, and *sfrp1a* expression in endothelial cells near Wnt9a-expressing somites suggests a potential regulatory interaction. Sfrps may also exhibit specificity: Sfrp1 interacts with canonical Wnt ligands like Wnt3a^61,62^, and non-canonical ligands like Wnt8^51^, whereas Sfrp2 modulates Wnt5a signaling^63^. These findings emphasize the role of Sfrps in both canonical and non-canonical pathways during HSPC development.

Understanding the role of Sfrp1a during HSPC development has important implications for hematopoietic malignancies. For example, low levels of Wnt signaling impair HSPC self-renewal and long-term maintenance, predisposing cells to acute myeloid leukemia, whereas intermediate levels sustain leukemic stem cells^15,64,65^. High Wnt signaling promotes HSPC exhaustion, clonal expansion, and transformation into T-cell acute lymphoblastic leukemia and acute myeloid leukemia^64,66,67,68,69,70,71^. These findings emphasize the importance of precise regulation of Wnt signaling for hematopoietic homeostasis. While mutations in Wnt ligands and Fzd receptors are rare in blood cancers, mutations in SFRPs, including SFRP1, are more common. Aberrant *SFRP1* expression is frequently observed in blood cancers^72,73,74^ (Fig. 6C), suggesting its potential as a biomarker for early detection and disease monitoring. Furthermore, engineering Sfrp-like molecules or small peptides mimicking their binding domains could enable localized and reversible modulation of Wnt signaling, offering a more precise therapeutic approach compared to broad spectrum Wnt inhibitors. Importantly, therapies targeting SFRP1 must carefully monitor dosage to balance therapeutic benefits and minimize side effects. Tissue-specific strategies targeting the spatial and temporal restriction of Sfrp expression, particularly in the bone marrow niche, could also enhance treatment options.

## METHODS

### Animals

Zebrafish were maintained and propagated according to Van Andel Institute and local Institutional Animal Care and Use Committee policies. AB* zebrafish were used as wild-type animals in all experiments. The *Tg(gata2b:KalTA4)^sd32Tg^* ^75^, *Tg(UAS:GFP)^mu2^*^71^ ^76^, *Tg(kdrl:NLS-mCherry)^y173Tg^* ^77^, *Tg(kdrl:Gal4)^bw9Tg^* ^55^*, Tg(7X TCF-X.laveis-siamois: eGFP)^ia4^* ^78^*, Tg(phldb1:KalTA4)^hzm7Et^* ^9^*, Tg(hsp:Gal4)^kca4Tg^* ^79^, *Tg(mpeg1:EGFP)^gl22Tg^* ^80^, *Tg(gata1:dsRed)^sd2Tg^* ^81^ lines have been previously described. The ATG MO for *sfrp1a* with the sequence 5’- GGACAAAGATGCAAGGGACTTCATT-3’, the splice-blocking MO for *sfrp1a* with the sequence 5’- AGTCATTTAGACTTACCGTTGGGTT-3’, and the scramble MO with the sequence 5’- CCTCTTACCTCAGTTACAATTTATA- 3’ were acquired from GeneTools. 1-cell stage zygotes were injected with 1 ng *sfrp1a* and 1 ng scramble MO. Embryos and larvae were cultured to the ages indicated in figures in Essential 3 (E3) medium (5 mM NaCl, 0.17 mM KCl, 0.33 mM CaCl2, 0.33 mM MgSO4, 10^-5^ % Methylene Blue).

Clustered regularly interspaced short palindromic repeats (CRISPR)–Cas9 was used to generate germline mutants for *sfrp1a*. We used a previously established method to selected two single guide RNAs based on their ability to cleave DNA *in vitro* as previously described^82^. Deletion of the proximal promoter and ATG start codon of *sfrp1a* was achieved using 100 ng *cas9* mRNA (Trilink) and 100 ng of each single guide RNA (5’- AAAGTCATGTTTTACATATC-3’ and 5’-TTGCATCTTTGTCCCTTTGG-3’). Primers used to validate the deletion and genotyping are available in Supplementary Table 1. Animals were generated with a 515 bp deletion. For simplicity, in the text, these are referred to as *sfrp1a^−/−^*. Mutations were confirmed by sequencing individual F1 animals. Zebrafish lines are available on request.

Sfrp1a (ENSDART00000051491.5) was PCR amplified from zebrafish cDNA. The transgenic plasmid for *UAS:Sfrp1a cmlc2:GFP* (referred to in the text as *UAS:Sfrp1a* for simplicity) was generated by inserting *sfrp1a* cDNA downstream of 4X tandem UAS in a construct with *cmlc2:gfp* and Tol2 recombination sites in the backbone, and sequence validated by full-plasmid sequencing. *Tg(UAS:Sfrp1a)* founders were established by injecting 25 pg of the *UAS:sfrp1a* generated plasmid with 100 pg transposase mRNA from the Tol2 kit at the one-cell stage^83^. The resultant animals were screened for GFP+ hearts and outcrossed to AB to establish germline founders.

The control and *sfrp1a* mRNA for experiments were synthesized using the SP6 mMessage machine kit (LifeTech), according to the manufacturer’s recommendations. The resultant mRNA was quantified by nanodrop, and dosages are indicated in figures and text. To assess potential off-target affects caused by mRNA injection alone, control mRNA was synthesized from an mCherry plasmid, which is expected to have no effect on development.

For heat shock experiments, fish were incubated at 35°C for 30 minutes and allowed to return to 28.5°C gradually. After an hour, 5 fish per biological condition were obtained for each heat shock timepoint and RNA was extracted for qPCR. At 40 hpf, zebrafish were isolated for WISH.

### Whole mount in situ hybridization (WISH)

Full length *sfrp1a* cDNA was ligated into pCRII-TOPO vector (Invitrogen) and digoxigenin (DIG)-labeled RNA probe was prepared by *in vitro* transcription with linearized construct based on manufacturer’s recommendations using the DIG-RNA labeling kit (Roche). Probes for *runx1*, *cmyb*, *kdrl*, *gata1a*, and *myod* and whole mount *in situ* hybridization (WISH) protocols have been previously described^6,9,84^.

### Fluorescently activated cell sorting and qPCR

Zebrafish were dissociated using Liberase TM (Roche) and filtered through a 80-µm filter as previously described^16^. Cells were sorted on a BD FACS Aria cell sorter and GFP+ cells were collected for qPCR. Total RNA was extracted from cells or embryos using a Quick-RNA Lysis Kit (Zymo Research, R1051) and cDNA was transcribed using iSCRIPT Reverse Transcription Supermix (BioRad, 1708841). qPCR was performed using PowerUp SYBR Green Master Mix (Thermo Fisher Scientific, A25777) according to the manufacturer’s recommendations and analyzed using the 2^-ΔΔCt^ method^85^. Primers used are available as a Supplementary Table 1.

### Confocal imaging and analysis

Live or fixed larval zebrafish were embedded in 2% UltraPure LMP Agarose (16520) in glass bottom dishes and covered with 1X E3 media. Imaging was performed using 25X immersion objective on the Andor Dragonfly 620-SR spinning disc mounted on a Leica DMi8 microscope equipped with 488, 561, and 640 nm laser lines.

For *7XTCF:eGFP; kdrl:mCherryNLS* imaging, Z-stacks were acquired using 0.49 μm step size and quantitative analysis of fluorescence intensity was performed using Imaris analysis software (10.2.0). The spot analysis tool was used to identify *7XTCF:GFP* and *kdrl:mCherryNLS* double positive cells along a 600 μm region of interest (ROI) of the floor of the dorsal aorta in all Z planes per image. Subsequent quantification of mean fluorescence intensity per identified cell was normalized to the total count of quantified cells. The brightness and contrast of all images were uniformly adjusted using histograms of intensity distributions to optimize visualization of fluorescence in representative images.

For *mpeg1:eGFP* and *gata1:dsRed* imaging, Z-stack images were acquired from 7 fields of view for fish imaged at 4 dpf and auto-stitched post-acquisition to encompass the whole animal. The stitched 3D images were cropped to include an ROI from the start of the yolk extension to end of caudal hematopoietic tissue. The ROI size was identical in all 4 dpf images. The surface model on Imaris software was used to detect the number of fluorescent cells.

## Acknowledgements

Imaging was performed in part in the Van Andel Institute Optical Imaging Core (RRID:SCR_021968). Research reported in this publication was supported by the National Institute of General Medical Science under Award Number R35GM142779 (SG), by the American Heart Association under Award Number 934891 (SG), National Institute of Cancer under Award Number T32CA251066 (ADI), and the American Cancer Society under Award Number PF-23-1037956-01-CCB (KAC) The content is solely the responsibility of the authors and does not necessarily represent the official views of the National Institutes of Health, American Heart Association, or American Cancer Society.

## Author Contributions

ADI, KC, ME, KO, KM, and CG designed, conducted and analyzed experiments. SG conceived, designed, supervised experiments and analysis. ADI and SG wrote the manuscript.

## Data and materials availability

All materials are available upon reasonable request.

---

**Supplementary Figure 1:**
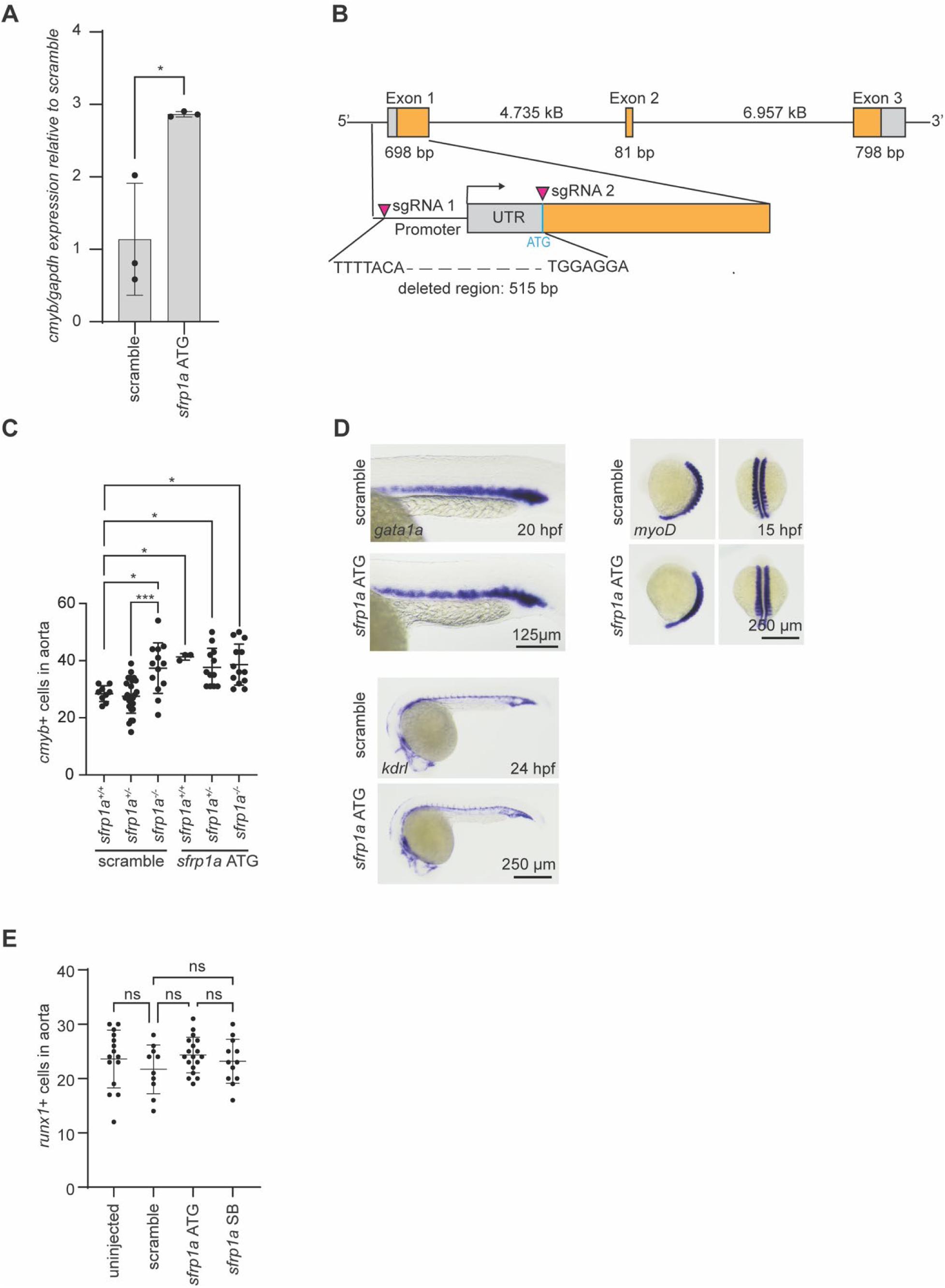
Sfrp1a loss of function does not affect other developmental processes. **A.** AB* embryos were injected with scramble morpholino and *sfrp1a* ATG morpholinos and analyzed by qPCR for *cmyb* at 40 hpf (*n=* 5 fish for each biological replicate, 3 biological replicates). Two-tailed Student’s t test, * p<0.05. **B.** *sfrp1a* mutants were generated by injection of two guide RNAs targeting upstream of the promoter and ATG start site on exon 1. **C.** *sfrp1a* mutants were injected with scramble or *sfrp1a* ATG morpholinos and expression of *cmyb* was analyzed by WISH at 40 hpf (*n*= 9, 26, 13, 3, 12, and 13 biological replicates from left to right). Each dot represents the mean number of cells in a biological replicate, the bars represent the mean and the error bars represent the standard deviation. One-Way ANOVA with post hoc Tukey comparisons, *p<0.05, **p<0.01, ***p<0.001. **D.** Scramble morpholino-injected or *sfrp1a* morpholino-injected embryos were analyzed for tissue-specific genes analyzed by WISH: representative images of primitive blood (*gata1a*), somites (*myod*), and vasculature (*kdrl*). **E.** AB* embryos were injected with scramble morpholino or *sfrp1a* morpholinos: ATG and splice blocking (SB), fixed at 40 hpf, and analyzed by WISH for *runx1* at 26 hpf (*n*= 15, 10, 18, and 12 biological replicates from left to right). Each dot represents a biological replicate, the bars represent the mean and the error bars represent the standard deviation. One-Way ANOVA with post hoc Tukey comparisons.

**Supplementary Figure 2.**
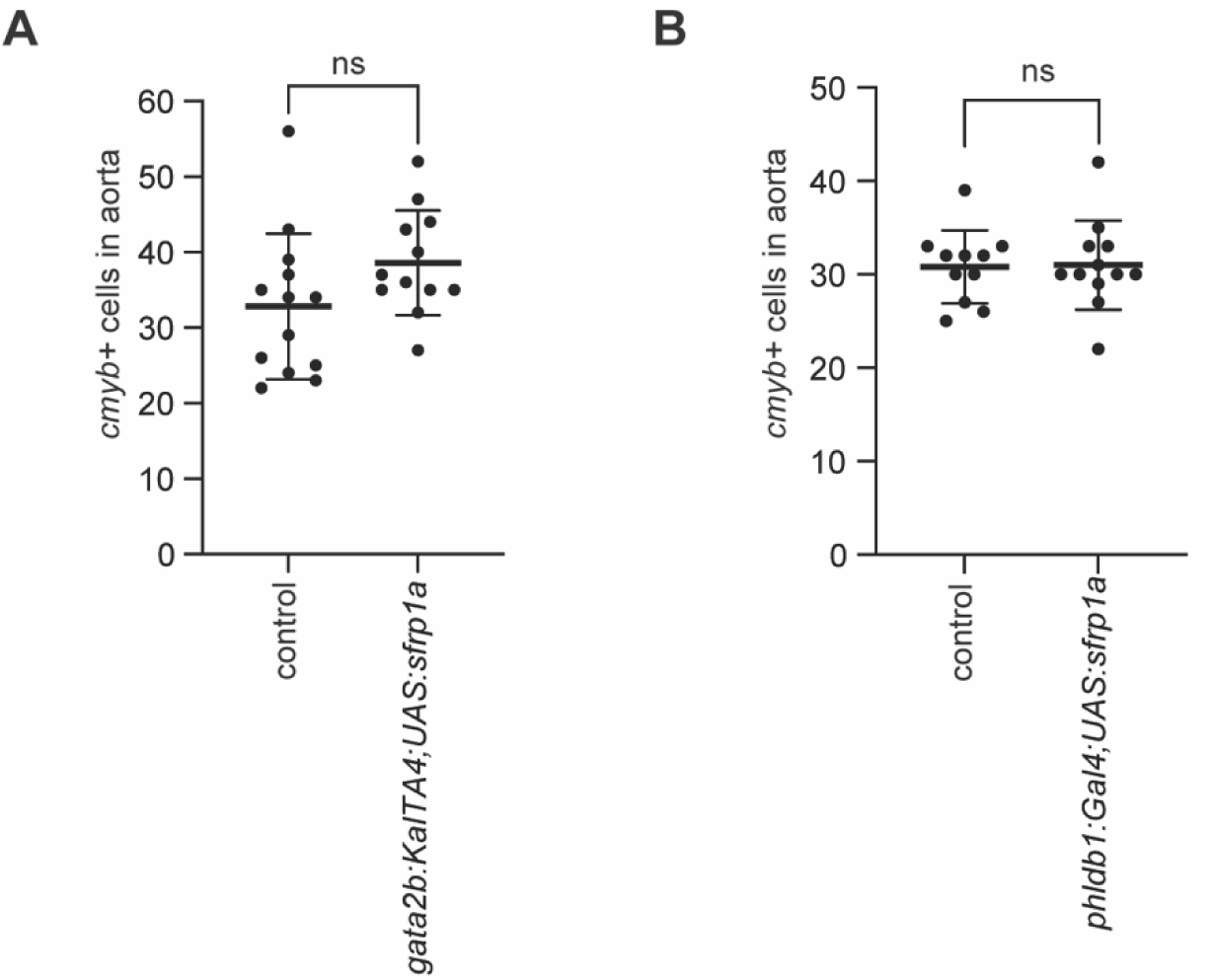
Sfrp1a expression in hemogenic endothelium or somites does not affect HSPC development. **A.** *gata2b:KalTA4* fish were crossed with *UAS:sfrp1a* fish, fixed at 40 hpf, and analyzed by WISH for *cmyb*. Control fish are wildtype siblings. Each dot represents a biological replicate (*n*= 13 and 12 biological replicates from left to right), the bars represent the mean and the error bars represent the standard deviation. Two-tailed Student’s t test. **B.** *phldb1:Gal4* fish were crossed with *UAS:sfrp1a* fish, fixed at 40 hpf, and analyzed by WISH for *cmyb*. Control fish are wildtype siblings. Each dot represents a biological replicate (*n*= 11 and 12 biological replicates from left to right), the bars represent the mean and the error bars represent the standard deviation. Two-tailed Student’s t test.

**Supplementary Figure 3.**
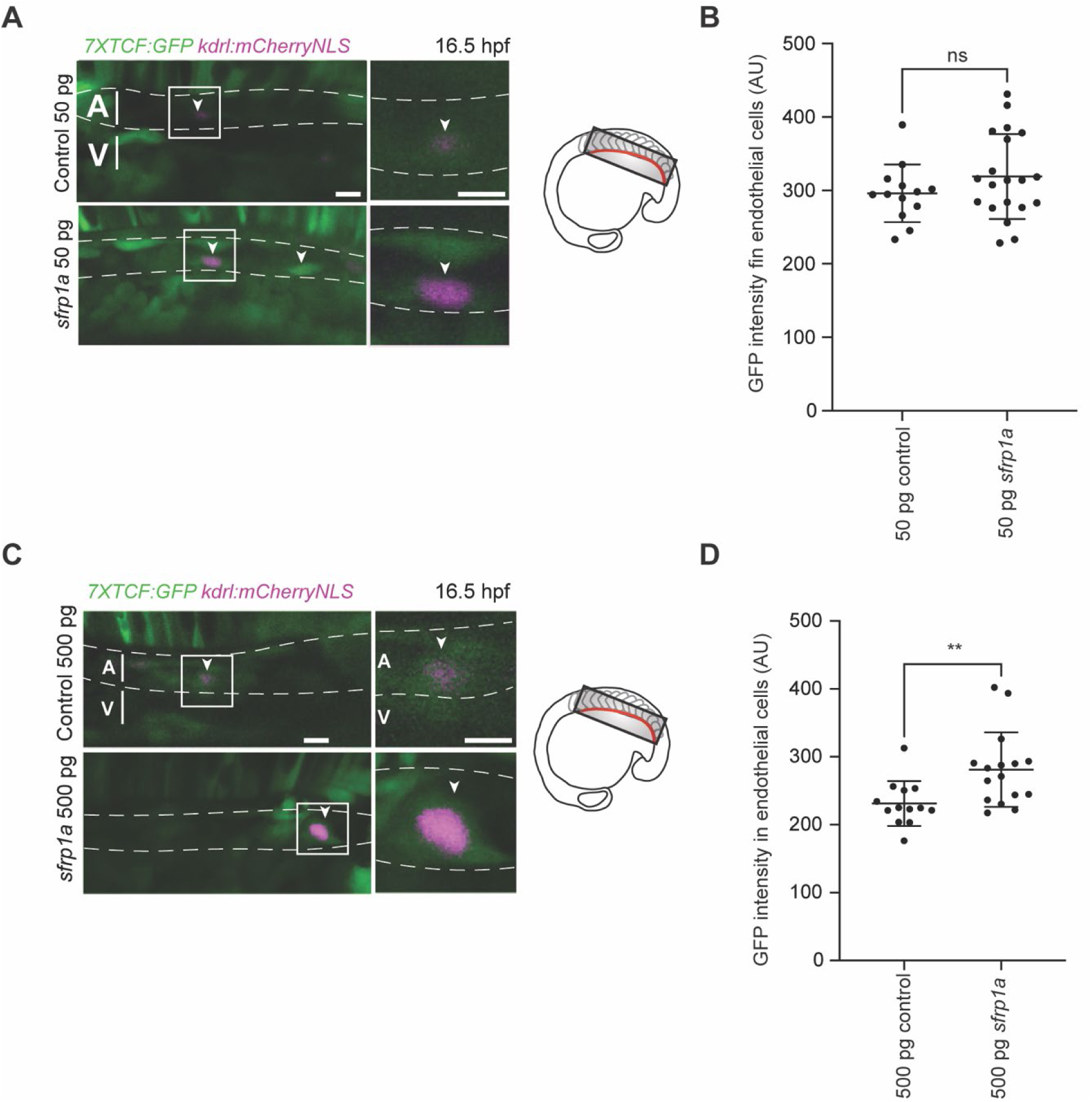
High *sfrp1a* overexpression leads to increased Wnt signaling in the developing endothelium. **A.** Representative images of *7XTCF:GFP; kdrl:mCherryNLS* fish injected with 50 pg control mRNA or 50 pg *sfrp1a* mRNA and the double positive cells were analyzed for GFP fluorescence at 16.5 hpf. Arrow heads point to double positive cells in the floor of the dorsal aorta. **B.** Quantification of GFP intensity in floor of the dorsal aorta (*n*= 13 and 20 cells in 8 and 9 fish, respectively). Mann-Whitney test. **C.** Representative images of *7XTCF:GFP; kdrl:mCherryNLS* fish injected with 500 pg control mRNA or 500 pg *sfrp1a* mRNA and the double positive cells were analyzed for GFP fluorescence at 16.5 hpf. Arrow heads point to double positive cells in the floor of the dorsal aorta. **D.** Quantification of GFP intensity in floor of the dorsal aorta (*n*= 13 and 16 cells in 7 fish). Mann-Whitney test, **p<0.01.

**Supplementary Figure 4.**
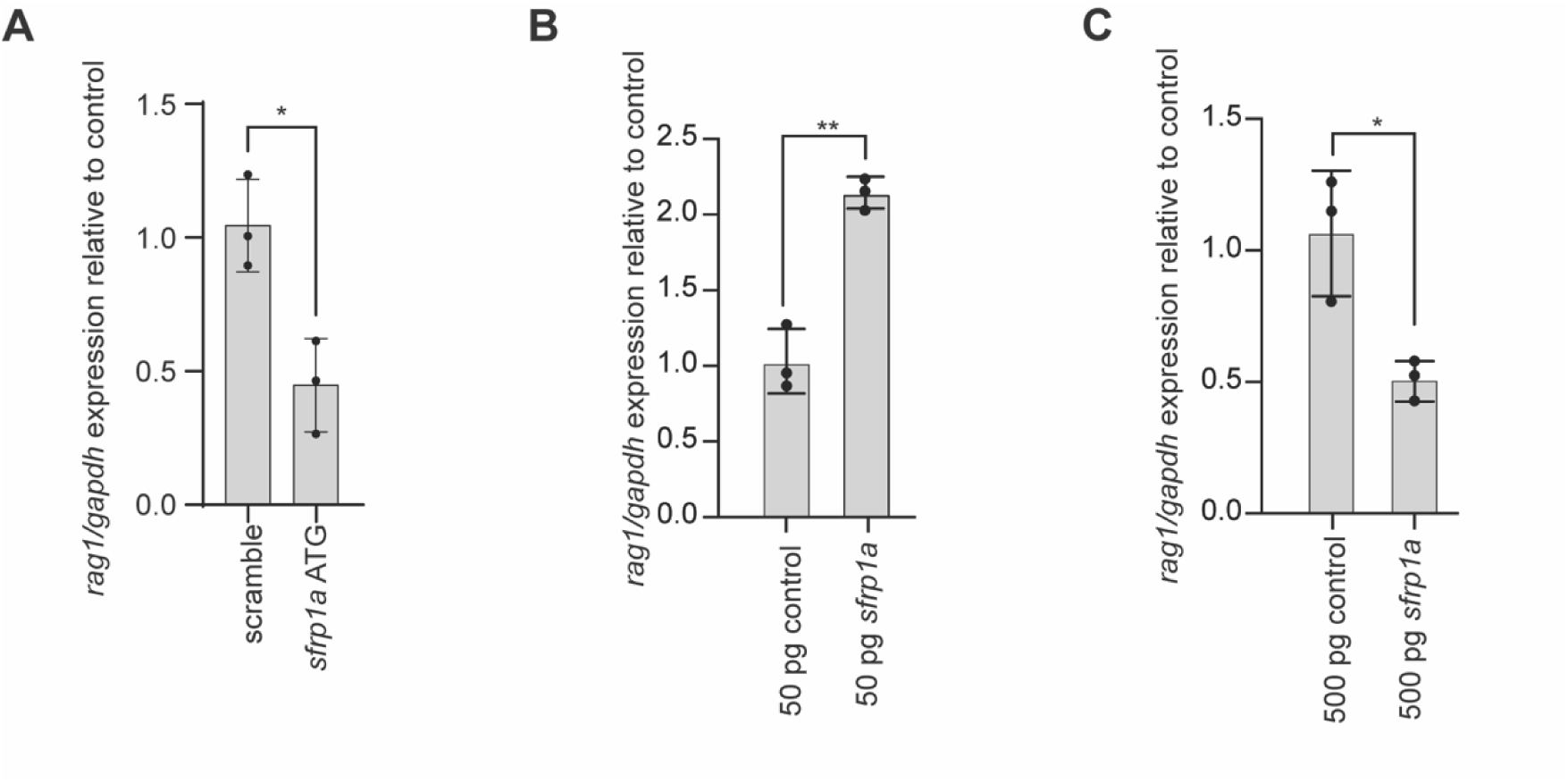
Sfrp1a impacts thymocyte expression in a dose-dependent manner. **A**. AB* embryos were injected with scramble and *sfrp1a* MO and expression of *rag1* was analyzed using qPCR (*n=* 5 fish for each biological replicate, 3 biological replicates). Two-tailed Student’s t test, *p<0.05. **B.** AB* embryos were injected with 50 pg control mRNA or 50 pg *sfrp1a* mRNA and expression of *rag1* was analyzed using qPCR (*n=* 5 fish for each biological replicate, 3 biological replicates). Two-tailed Student’s t test, **p<0.01. **C.** AB* embryos were injected with 500 pg control mRNA or 500 pg *sfrp1a* mRNA and expression of *rag1* was analyzed using qPCR (*n=* 5 fish for each biological replicate, 3 biological replicates). Two-tailed Student’s t test, *p<0.05.

**Supplementary Figure 5:**
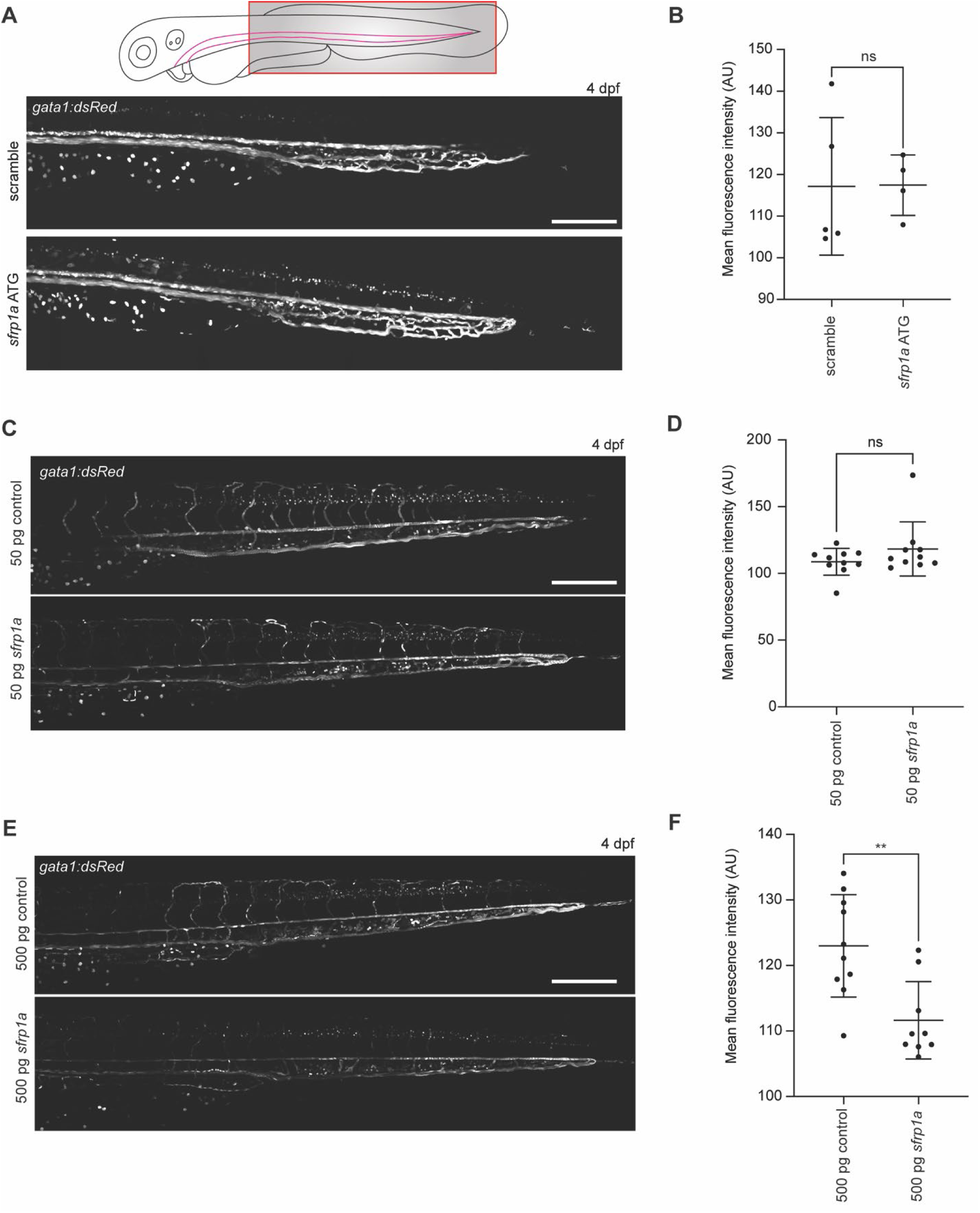
Sfrp1a does not affect erythromyeloid differentiation in a dose-dependent manner. **A.** Representative confocal Z-stacks of *gata1:dsRed* fish injected with scramble or *sfrp1a* ATG morpholino, embedded in agar at 4 dpf and imaged at 25X. Scale bar = 200 µm. **B**. Quantification of **A** marked by dsRed+ cells (*n*= 5 and 4 biological replicates from left to right). Each dot represents the mean number of cells in a biological replicate, the bars represent the mean and the error bars represent the standard deviation. Mann-Whitney test. **C.** Representative confocal Z-stacks of *gata1:dsRed* fish injected with 50 pg control mRNA or 50pg *sfrp1a* mRNA, embedded in agar at 4 dpf and imaged at 25X. Scale bar = 200 µm. **D**. Quantification of **C** marked by dsRed+ cells (*n*= 10 biological replicates). Each dot represents the mean number of cells in a biological replicate, the bars represent the mean and the error bars represent the standard deviation. Mann-Whitney test. **E.** Representative confocal Z-stacks of *gata1:dsRed* fish injected with 500 pg control mRNA or 500 pg *sfrp1a* mRNA, embedded in agar at 4 dpf and imaged at 25X. Scale bar = 200 µm. **F**. Quantification of **E** marked by dsRed+ cells (*n*= 10 and 9 biological replicates from left to right). Each dot represents the mean number of cells in a biological replicate, the bars represent the mean and the error bars represent the standard deviation. Mann-Whitney test, **p<0.01.

**Supplementary Table 1:**
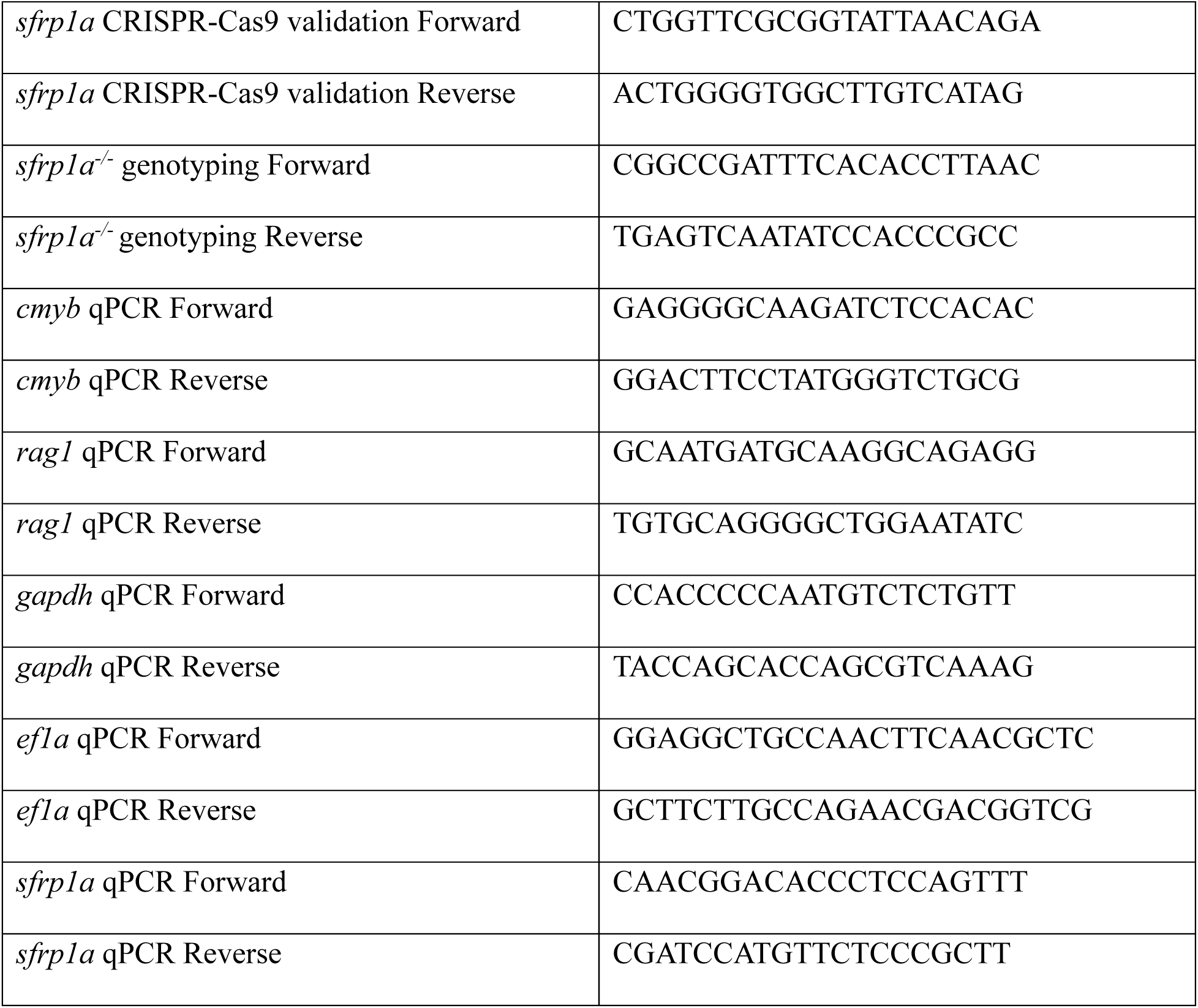
PCR and qPCR primers used.

